# A comparative investigation of the mannose binding interface in DC-SIGN and MRC1 carbohydrate recognition domains with all-atom molecular dynamics simulations

**DOI:** 10.64898/2026.01.16.699853

**Authors:** Sina Geissler, Sophie Sacquin-Mora

## Abstract

Protein-carbohydrate interactions play a key role in numerous biological processes, including immune response, and glycan-based ligands that can target specific protein receptors on a cell surface represent promising candidates for therapeutics applications. For example, in retinoblastoma, the DC-SIGN mannose receptor is overexpressed on the surface of pathogenic cells and represents an interesting target for mannose-based ligands. At the same time, these ligands should not bind to the MRC1 receptor, which is expressed by adjacent, healthy, retinal pigment epithelial cells and presents a carbohydrate recognition domain (CRD) similar to the one of DC-SIGN. Therefore, the challenge remains to obtain a detailed picture of the recognition process between carbohydrates and proteins, in order to design effective and selective therapeutic compounds. In this work we used classical, all-atom molecular dynamics (MD) simulations to investigate the interaction between several mannose based ligands and the CRDs from DC-SIGN and MRC1. The analysis of the protein-carbohydrate contacts from the resulting trajectories highlights the variability of the mannose binding modes on both CRDs, and shows how the increased affinity of mannose for the MRC1 CRD can be related to a specific mannose binding state that is not accessible in the DC-SIGN CRD.

**TOC image:** For Table of Contents use only

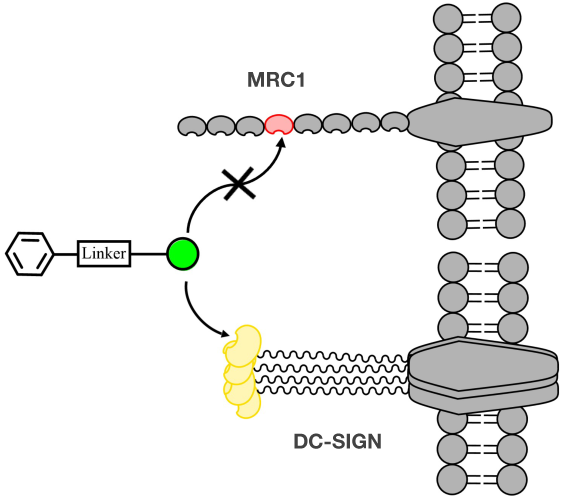

## 1. Introduction

Carbohydrates, or glycans, represent the most abundant class of organic molecules on Earth,^1^ and are a central component of life, both as a primary energy source^2^ and as key player of cellular functions such as signaling and recognition.^3, 4^ Glycan diversity covers a broad range of size, shape and functions,^5, 6^ which is mirrored by the diversity of protein-carbohydrate interactions.^7–9^ These interactions can be related to an organism’s immune response,^10, 11^ but also to numerous diseases, such as cancer,^12, 13^ and host-pathogen recognition.^14–17^ As a consequence, glycans, with their ability to bind specific protein targets, appear as promising candidates for therapeutic applications.^1, 18–21^ One example of such biomedical applications is the development of targeted photodynamic therapies (PDTs), which use photosensitizers (PSs) where a porphyrin group is connected to one or several mannose groups, and are very efficient against small solid tumors accessible to light.^22, 23^

Mannose receptors (MRs) are transmembrane proteins and a subgroup of the C-type lectins superfamily. This large family of carbohydrate binding, non-enzymatic, non-immunoglobulin proteins, binds glycans in a Ca^2+^ dependent manner via their C-type lectin-like domain (CTLD), which is also termed as carbohydrate recognition domain (CRD).^24–28^ While these proteins can display a wide range of quaternary structures (see Figure 1a), their CRDs present a high structural similarity. The canonical fold comprises 110-140 residues, and consists of six to seven β-strands organized in two β-sheets and flanked by two ⍺-helices (see Figure 2b).

**Figure 1:**
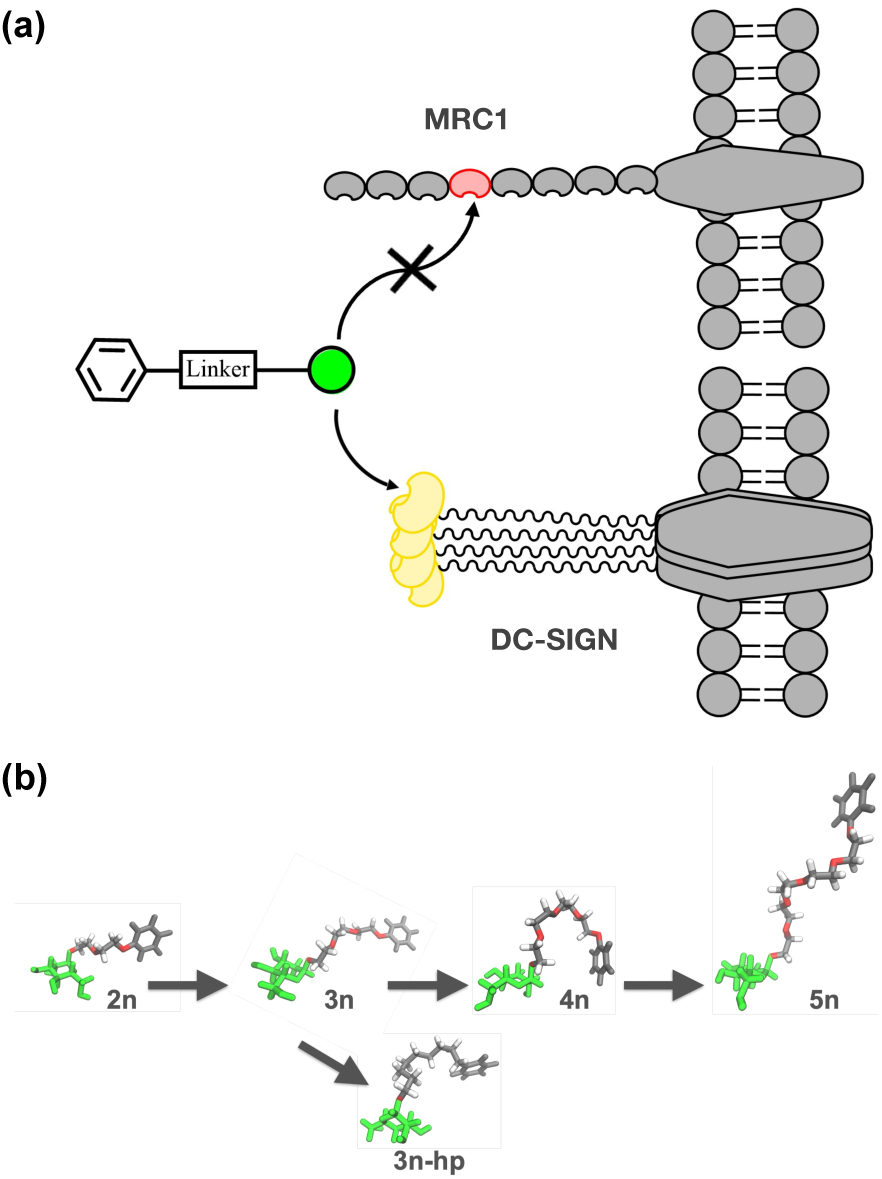
(a) Schematic representations of MRC1 and DC-SIGN receptors with the mannose-based ligand (the mannose moiety is shown in green), the modeled carbohydrate recognition domains have been highlighted in red and yellow for MRC1 and DC-SIGN respectively. (b) Graphic summary of the mannose based ligands tested in the simulations. For ligands 2n-5n, the linker between the mannose and benzene ring comprises 2 to 5 (-CH_2_-CH_2_-O-) units. For the more hydrophobic 3n-hp ligand (lower line) the linker between the mannose and benzene ring comprises 3 (-CH_2_-CH_2_-CH_2_) units. Figures 1b, 2b-d, 4b, 6bc, 8, 9a and 10 were prepared using Visual Molecular Dynamics (VMD).^57^

**Figure 2:**
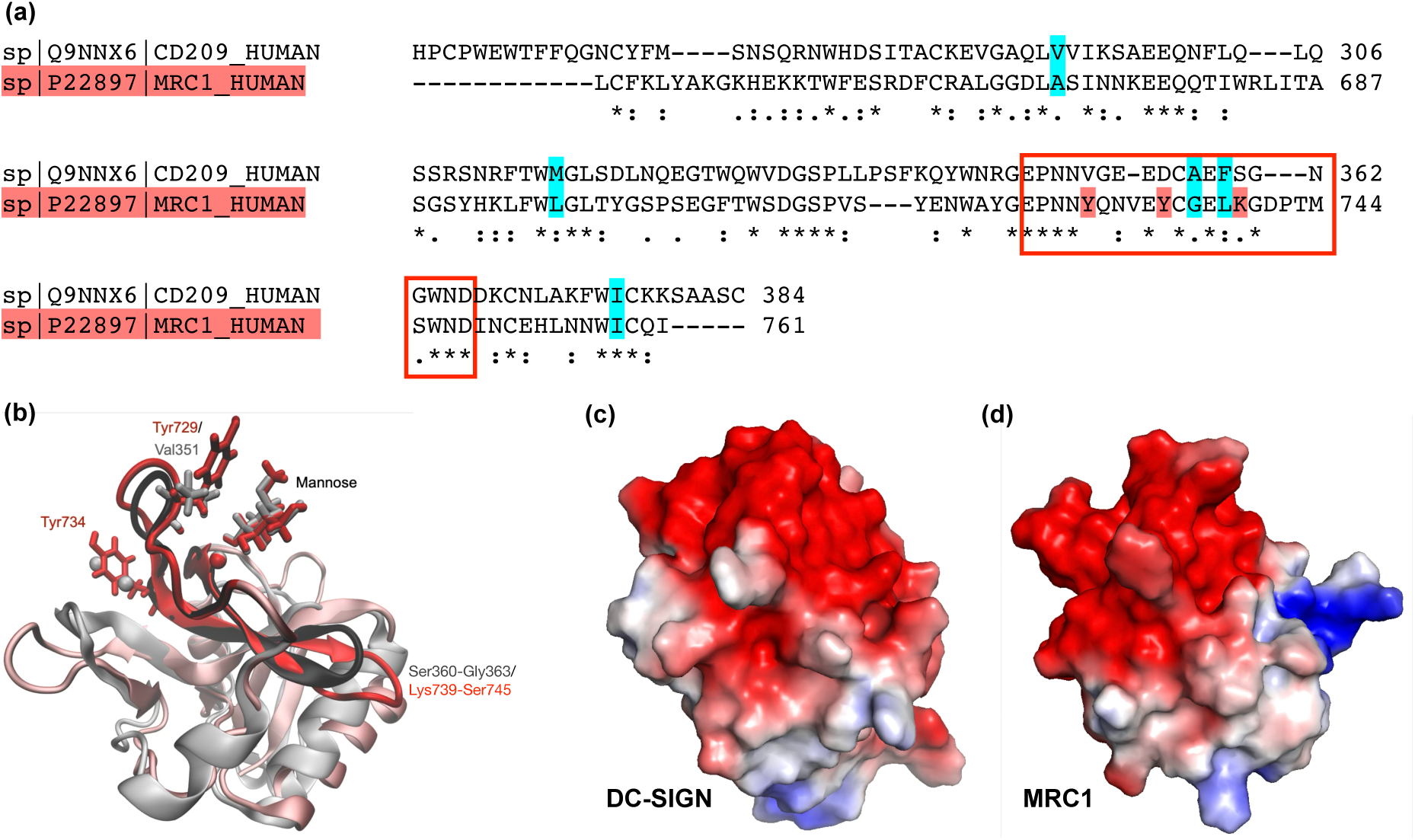
(a) Sequence alignement for the CRDs of DC-SIGN (upper line) and MRC1 (lower line). The red rectangle highlights the mannose binding pocket, the conserve rigid residues (also shown on Figure 3cd) are highlighted in cyan. (b) Aligned crystallographic structures for the DC-SIGN (in gray) and MRC1 (in red) CRDs. Surface electrostatic potentials for the CRDs, positive potentials are colored in blue and negative potentials in red (c) DC-SIGN, (d) MRC1.

MRs, can be overexpressed on the surface of pathogenic cells,^24, 29^ and therefore represent a potential target for PDTs. This is the case of the dendritic cell-specific ICAM-3-grabbing non-integrin, abbreviated as DC-SIGN, a protein from dendritic cells which plays a central role in immunity.^30^ The extracellular domain of DC-SIGN comprises a neck region ending in the CRD, and assembles in a coiled-coil homotetramer via the neck domain (see Figure 1a).^31^ Retinoblastoma (Rb) is a cancer which develops at the level of photoreceptors in the retina,^32^ and Rb cells overexpress DC-SIGN, thus making this MR a potential target for mannosylated porphyrins.^33^

However, the efficiency of a PDT treatment relies both on the PS’s affinity for its target, but also on its specificity. In the present case, this would require the mannosylated ligands to not bind the human C-type 1 mannose receptors (MRC1) that are expressed on the surface of the healthy, adjacent, retinal pigment epithelium cells.^34, 35^ Unlike DC-SIGN, the extracellular part of MRC1 comprises a single chain with eight successive C-type lectin domains, where only the fourth one (CRD4) can bind monosaccharide ligands on its own (see Figure 1a).^36^ The CRD from DC-SIGN and CRD4 from MRC1 both present similar folds and sequences. As a consequence, the design of adequate mannose-based ligands for PDT will require to characterize in detail the interface between mannose and the CRD’s glycan binding site.

Numerous experimental methods are currently available for the characterization of the structure and affinity of protein-carbohydrate interfaces,^37^ including (but not limited to) X-ray crystallography,^38, 39^ NMR spectroscopy,^40^ Cryo-EM^41^ and isothermal titration calorimetry (ITC).^42^ The high flexibility of glycans and the low-affinity of some carbohydrate recognition mechanisms^27^ can however represent a challenge for experimental approaches, and computational analyses can help improving our understanding of these interactions.^43–46^ In this context, we used classical, all-atom molecular dynamics (MD) simulations to investigate the interaction between several mannose based ligands and the CRDs from DC-SIGN and MRC1 on the atomic level. The analysis of the protein-carbohydrate contacts from the resulting trajectories highlights the variability of the mannose binding modes on both CRDs, and shows how the increased affinity of mannose for the MRC1 CRD can be related to a specific mannose binding state that is not accessible in the DC-SIGN CRD.

## 2. Material and Methods

### Starting models and MD simulations

For the DC-SIGN-CRD, we used the Protein Data Bank (PDB)^47^ crystal structure 2XR5^48^ (DC-SIGN with mannose, 1.42 Å resolution) as the starting conformation. Simulations of the MRC1-CRD were based on the 7JUB crystal structure^49^ of the CRD4 of MCR1 (MRC1 with methyl-mannoside, 1.20 Å resolution). For this system, methyl-mannoside was changed to mannose using the CHARMM-GUI^50^ web server (https://www.charmm-gui.org/). In addition to single mannose, five mannose-based ligands were also modeled and parameterized on CHARMM-GUI,^51^ using NMR-experimental structures provided by our collaborators as first input (see Figure 1b). The ligands were then superposed to the mannose group in the crystal structures using Pymol^52^ (Version 3.1.6.1) to generate a starting conformation of the ligand bound to the receptor. All starting structures were solvated with TIP3P water^53^ using the CHARMM-GUI server. The calcium cations initial positions (three for DC-SIGN and only one for MRC1) were taken from the crystal structures as they are important for the ligand binding. Potassium and chloride ions were added to the solution with a physiological concentration of 0.15 mol.L^-1^. The pH of the system was set to 7 and the histidine residues were singly deprotonated to form neutral histidines. For ligands bigger than a simple mannose, the buffer distance around the protein was set to 15 Å (10 Å for simulations with only mannose) to define the rectangular box dimension. The difference in length and volume between the DC-SIGN CRD (131 residues, more elongated) and the MRC1 CRD (122 residues, more compact shape) and the different sizes of the ligands and their All-atom molecular dynamics simulations were carried out using GROMACS(v.2023.2),^54^ and the CHARMM36m^55^ force-field (see the force-field discussion in the supplementary information), with a time step of 2 fs. All simulations were done under periodic boundary conditions with fast smooth Particle-Mesh Ewald electrostatics.^56^ First, energy minimization was done in 5000 steps with the steepest decent algorithm and a Verlet cutoff scheme. The cutoff is set by the Verlet-buffer-tolerance at its default value of 0.005 kJ.mol^-1^.ps^-1^. The positions for the solutes backbone atoms were restrained with a force constant value of 400 kJ.mol.nm^-2^ and side chain atoms with a force constant of 40 kJ.mol.nm^-2^. H-bonds were constrained with the LINCS^57^ algorithm. The equilibration step was done in six simulations with decreasing position and dihedral restraints ( with backbone force constants of 4000-2000-1000-500-200-50 kJ.mol^-1^.nm^-2^, side chain force constants of 2000-1000-500-200-50-0 kJ.mol^-1^ .nm^-2^ and dihedral force constants of 1000-400-200-200-100-0 kJ.mol^-1^.nm^-2^ in each simulation respectively), the ‘md’ integrator (leap-frog algorithm), and the Verlet cutoff scheme set again by the default Verlet-buffer-tolerance, and 125000 steps for the first three equilibration runs (corresponding to 0.125 ns per simulation) and 250000 steps for the last three equilibration runs (corresponding to 0.5 ns per simulation). Pressure and temperature coupling was done using the Berendsen^58^ algorithm. During the 500 ns of the production step, pressure coupling to 1.0 Bar was done using the Parrinello-Rahman algorithm,^59^ and we used the Nose-Hoover algorithm^60^ for temperature coupling to room temperature (295.15 K). The root mean square deviation (RMSD) plot showing the equilibration of the protein structure during the whole procedure is available in Figure S1. For each ligand-receptor pair, we ran three replicas for the 500 ns production steps in the NPT ensemble, and frames were saved every 100 ps during production. All MD parameter files are available online (https://github.com/sinageissler/DC-MRC1-paper1), and Table 1 summarizes all the systems that were simulated during this study.

**Table 1:** Summary of the MD simulations performed in the study.

| | Ligand atoms | Protein atoms/<br>residues | K<sup>+</sup> and Cl<sup>-</sup> ions | Water molecules | Box size [$\text{\AA}^3$] |
| --- | --- | --- | --- | --- | --- |
| DC | - | 2037/131 | 23, 27 | 8181 | 64<sup>3</sup> |
| man-DC | 24 | 2037/131 | 27, 31 | 9417 | 67<sup>3</sup> |
| 2n-DC | 48 | 2037/131 | 38, 42 | 13573 | 75x75x74 |
| 3n-DC | 55 | 2037/131 | 52, 56 | 18420 | 82x82x83 |
| 3nhp-DC | 61 | 2037/131 | 56, 60 | 19879 | 85x85x84 |
| 4n-DC | 62 | 2037/131 | 38, 42 | 13548 | 75<sup>3</sup> |
| 5n-DC | 69 | 2037/131 | 47, 51 | 16563 | 80<sup>3</sup> |
| MRC | - | 1920/122 | 21, 19 | 6748 | 60<sup>3</sup> |
| man-MRC | 24 | 1920/122 | 21, 19 | 6736 | 61x61x59 |
| 2n-MRC | 48 | 1920/122 | 34, 32 | 11394 | 71<sup>3</sup> |
| 3n-MRC | 55 | 1920/122 | 34, 32 | 11380 | 71<sup>3</sup> |
| 3nhp-MRC | 61 | 1920/122 | 31, 29 | 10414 | 69<sup>3</sup> |
| 4n-MRC | 62 | 1920/122 | 33, 31 | 10971 | 70<sup>3</sup> |
| 5n-MRC | 69 | 1920/122 | 33, 31 | 10982 | 71x71x69 |

### Analysis of the MD Trajectories

A sequence alignment of the two receptor pocket sequences was done on the UniProt server,^61^ surface electrostatics were calculated on the APBS server (https://server.poissonboltzmann.org/)^62^ using the crystal structure PDB files as input. Unbinding of a ligand was defined via the Colvars dashboard in VMD.^63, 64^ The collective variable was defined as a distance between the pocket residues and the ligand, the ligand was defined as bound until the last full nanosecond before the collective variable surpassed the value of 10 Å. Molecular mechanics/Poisson-Boltzmann surface area (MM/PBSA)^65^ binding enthalpies between the ligand and the CRD for full trajectories were calculated using the GROMACS tool and all the simulated frames with the ligand bound across all three replicas of each simulation as input data, for better sampling of the bound positions. All the analysis scripts are available online (https://github.com/sinageissler/DC-MRC1-paper1)

RMSD, root mean square fluctuations (RMSF) values, and covariance matrices on the C⍺ coordinates were calculated with GROMACS,^54^ while MDAnalysis^66, 67^ was used for calculating interatomic distances. The different ligand binding states were defined based on characteristic distances between the OH-groups of the mannose moiety and specific residues of the CRDs binding pocket. The cutoff distance was set to 3.5 Å, which roughly corresponds to the length of hydrogen bonding. Frames with receptor-ligand conformations corresponding to the same binding state were then binned together into a single trajectory, which was used to calculate the contact maps shown in Figures 6 and 8. The binding affinity for each specific mannose binding state was calculated using the MM/PBSA method for states with more than 100 frames only, to ensure sufficient sampling.

### Coarse-grain Brownian Dynamics simulations

#### Rigidity profile of a protein

Coarse-grain Brownian Dynamics (BD) simulations were run using the ProPHet (Probing Protein Heterogeneity, available online at https://bioserv.rpbs.univ-paris-diderot.fr/services/ProPHet/) program^68–70^. In this approach, the protein is represented using an elastic network model. Unlike most common coarse-grain models where each residue is described by a single pseudoatom,^71^ ProPHet uses a more detailed representation^72^ that involves up to 3 pseudoatoms per residue and enables different amino acids to be distinguished. Pseudoatoms closer than the cutoff parameter *R_c_* = 9 Å are joined by Gaussian springs which all have identical spring constants of γ_struct_ = 0.42 N.m^-1^ (0.6 kcal.mol^-1.^Å^-2^). The springs are taken to be relaxed for the initial conformation of the protein. The simulations use an implicit solvent representation via the diffusion and random displacement terms in the equation of motion,^73^ and hydrodynamic interactions are included through the diffusion tensor.^74^

Mechanical properties are obtained from 200,000 BD steps at an interval of 10 fs and a temperature of 300 K. The simulations lead to deformations of roughly 1.5 Å root-mean-square deviation with respect to the protein starting conformation (which, by construction, corresponds to the system’s equilibrium state). The trajectories are analyzed in terms of the fluctuations of the mean distance between each pseudoatom belonging to a given amino acid and the pseudoatoms belonging to the remaining residues of the protein. The inverse of these fluctuations yields an effective force constant *k_i_* describing the ease of moving a pseudoatom with respect to the overall protein structure.

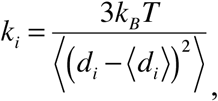

where 〈〉 denotes an average taken over the whole simulation and *d_i_=* 〈*d_ij_*〉*_j*_* is the average distance from particle *i* to the other particles *j* in the protein (the sum over *j\** implies the exclusion of the pseudoatoms belonging to residue *i*). The distance between the C_α_ pseudoatom of residue *i* and the C_α_ pseudoatoms of the adjacent residues *i-1* and *i+1* are excluded since the corresponding distances are virtually constant. The force constant for each residue *k* is the average of the force constants for all its constituent pseudo atoms *i*. We will use the term *rigidity profile* to describe the ordered set of force constants for all the residues of the protein.

## 3. Results and discussion

### Structural similarity of the DC-SIGN and MRC1 glycan binding sites

Despite a low sequence identity (around 32%), both CRDs present very similar crystallographic structures (1.3 Å rmsd on the heavy atoms), with a set of five conserved residues lining the Ca^2+^ cation involved in glycan binding, namely Glu347/725, Asn349/727, Glu354/733, Asn365/747 and Asp366/748 for DC-SIGN and MRC1 respectively (see the sequence alignment in Figure 2a). The most notable differences are the presence of two tyrosine residues in MRC1 (highlighted in red in Figure 2a). Tyr729 lies in the glycan binding pocket (instead of a valine for DC-SIGN), and Tyr734 lies in the pocket occupied by two additional Ca^2+^ ions in DC-SIGN. In addition, the Lys739-Ser745 loop is slightly longer for MRC1 than its Ser360-Gly363 DC-SIGN counterpart, leading to a deeper groove for the glycan binding pocket (See Figure 2b). The Lys739 residue at the beginning of the loop, instead of Ser360 for DC-SIGN, leads to the presence of a positive patch on the CRD’s surface for MRC1. The RMSD evolution along the MD trajectories for the CRDs with mannose shows stable structures, with the RMSD increase along time for the DC-SIGN replicas being due to the mobility of the N-terminal tail (see Figure 3a, upper panel). The RMSD evolution for the glycan binding pocket only suggests that it is less flexible for MRC1 (see Figure 3a, lower panel) and this concurs with the RMSF profiles shown in Figure 3b. Covariance analysis on the trajectories for the CRDs in their apo state shows that the first five eigenvectors for DC-SIGN are mostly localised on the N-terminus and on the glycan binding pocket, while for MRC1, the dynamics are more broadly distributed over the whole protein sequence (see Figure S2). Finally, for both CRDs, the rigidity profile resulting from the coarse-grain simulations highlight a well-defined, conserved, rigid hydrophobic core (see Figure 4), with the DC-SIGN CRD being slightly more rigid on average. One should note that, since protein mechanical properties are heterogeneously distributed over the whole protein structure,^69^ the CRD from DC-SIGN can simultaneously have a more rigid core and a more flexible glycan binding site than the CRD from MRC1.

**Figure 3:**
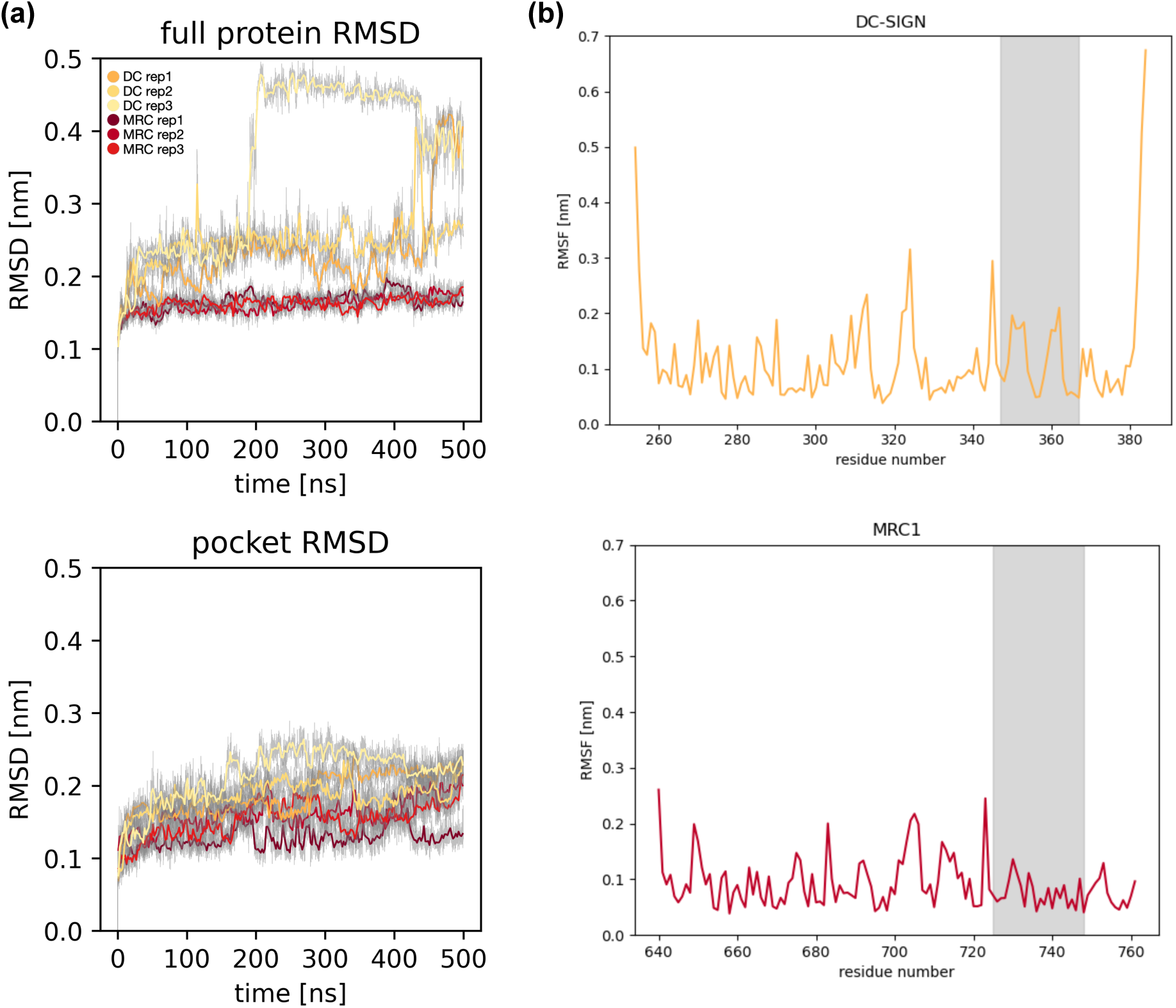
RMSD as a function of time for the DC-SIGN (yellow lines, three replicas) and MRC1 (red lines, three replicas) CRDs with mannose. Upper panel: full protein, lower panel: glycan binding pocket only. For clarity, all colored curves were smoothed using a 20-frames sliding window. (b) RMSF for the apo-protein (with calcium ions and without ligands) on the three concatenated replicas, the glycan biding pocket location is highlighted in grey. Upper panel: DC-SIGN, lower panel: MRC1.

**Figure 4:**
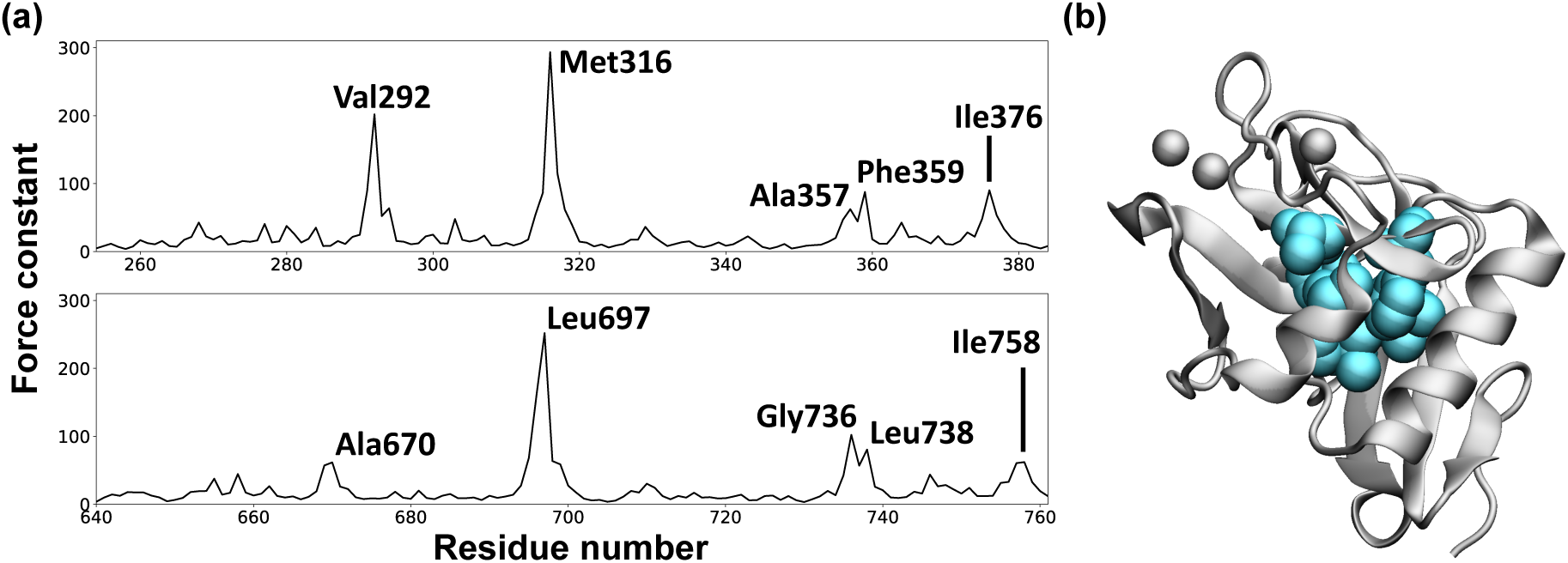
(a) Force constants (in mol^-1.^Å^-2^) for the DC-SIGN (upper line) and MRC1 (lower line) CRDs. (b) Crystallographic structure of the DC-SIGN CRD with the rigid residues annotated in panel-a highlighted in cyan.

### Comparing the DC-SIGN and MRC1 binding affinities for the different ligands

A striking result from the all-atom MD simulations is the instability of the mannose/ CRD interface for DC-SIGN, with the systematic unbinding of the ligand within the first 100 ns of each trajectory for all the replicas and all the tested ligands (see Figure 5). The unbinding events for each replica are also visible in the ligand RMSD plots shown in Figure S3, as they result in large RMSD fluctuations. This, however, is in line with our current knowledge of DC-SIGN-carbohydrate interactions.^75, 76^ In the case of MRC1, the ligand/CRD interaction appears to be more stable, especially for the 3n and 4n ligands, which manage to stay bound in the glycan binding pocket for the complete duration of the simulation for at least one replica. The difference in stability for mannose bound on DC-SIGN and MRC1 can be related to the fluctuations in the distance between the glutamate residues surrounding the Ca^2+^ ion, (Glu347/725 and Glu354/733, see Figure6b-c), which are essential for mannose binding in the crystallographic state. While this distance remains stable for MRC1, it fluctuates much more in DC-SIGN (see Figure S4).

**Figure 5:**
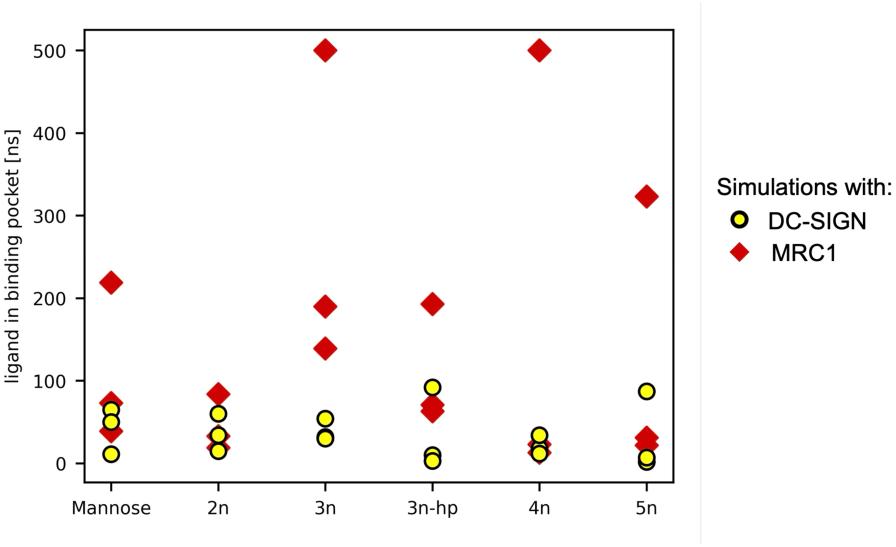
Ligand residency time in the glycan binding pocket for all possible ligands/CRD combinations (6*2*3 replicas).

For MRC1, the ligands linker section (which connects the mannose moiety to the benzene ring, see Figure 1b) is in contact with the polar Tys729 (see Figure S5), which might contribute to stabilizing the CRD/ligand interface. Hence, for the 3n case, we tested whether a more hydrophobic linker (with no oxygen atoms) could improve the ligand/DC-SIGN interaction while simultaneously disrupting the ligand/MRC1 interaction. The resulting 3n-hp ligand shows no significant increase of its affinity for DC-SIGN, while being slightly more unstable than the 3n ligand when interacting with MRC1. All the resulting ligand/CRD binding affinities are available in Table 2 and the values are of the same order of magnitude as those obtained in earlier simulation works.^77, 78^

**Table 2:**
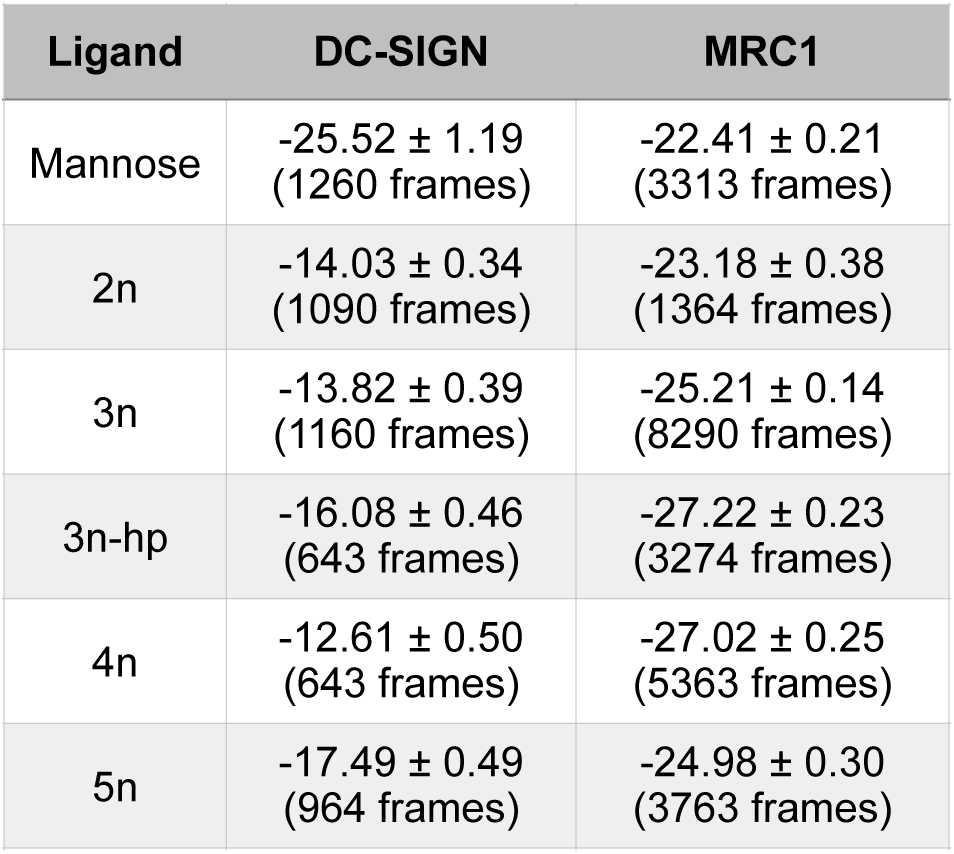
Average Ligand/CRD binding enthalpies (in kcal.mol^-1^)

| Ligand | DC-SIGN | MRC1 |
| --- | --- | --- |
| Mannose | -25.52 ± 1.19<br>(1260 frames) | -22.41 ± 0.21<br>(3313 frames) |
| 2n | -14.03 ± 0.34<br>(1090 frames) | -23.18 ± 0.38<br>(1364 frames) |
| 3n | -13.82 ± 0.39<br>(1160 frames) | -25.21 ± 0.14<br>(8290 frames) |
| 3n-hp | -16.08 ± 0.46<br>(643 frames) | -27.22 ± 0.23<br>(3274 frames) |
| 4n | -12.61 ± 0.50<br>(643 frames) | -27.02 ± 0.25<br>(5363 frames) |
| 5n | -17.49 ± 0.49<br>(964 frames) | -24.98 ± 0.30<br>(3763 frames) |

### Investigating the mannose binding modes in the glycan binding pockets on DC-SIGN and MRC1

The mannose binding mode in the CRDs crystallographic structure has already been described in the experimental literature,^48, 49^ with the mannose OH3 and OH4 groups forming contacts with the calcium ion from the glycan binding pocket. In addition, the OH2 and OH3 groups interact with Glu354/Glu733 (highlighted in red on Figure 6b,c), while OH4 interact with Glu 347/Glu 725 (highlighted in blue on Figure 6b,c) for DC-SIGN and MRC1 respectively.

**Figure 6:**
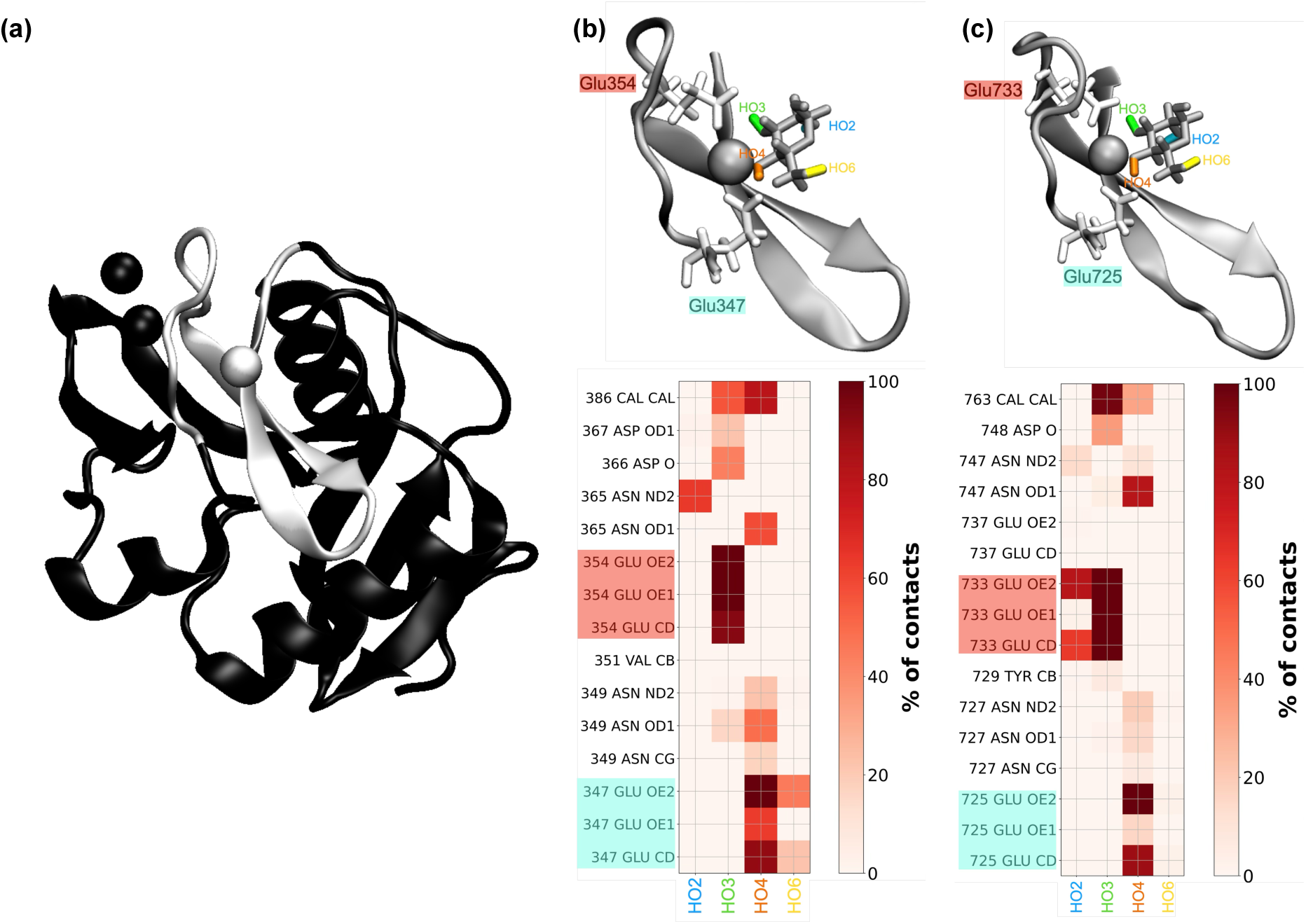
(a) Crystallographic structure of the DC-SIGN CRD with the glycan binding site highlighted in white. Mannose binding mode in the crystallographic structures with the corresponding contact map (b) DC-SIGN, (c) MRC1. The blue/green/orange/yellow color code to distinguish the mannose OH2/OH3/OH4/OH6 groups is kept in Figure 8.

However, for both receptors, and all ligands, the mannose group leaves its crystallographic initial position at an early stage of the simulation and explores additional binding modes in the CRDs glycan binding pocket that are summarized in Figure 7. In *state A*, which was observed for both the DC-SIGN and MRC1 CRDs, the mannose moiety rotates so that its OH2 interacts with Glu347/725, while both OH3 and OH4 form contacts with Glu354/733 (see Figure 8a,c). This binding mode can be related to the minor mannose binding mode described by Cramer,^31^ and was also observed by Feinberg et al.^49^ for fucose binding to the MRC1 CRD. Two other binding modes were observed for mannose in the CRDs glycan binding pockets. *State B* is also common to the DC-SIGN and MRC1 CRDs and sees OH3 form contacts with the Ca^2+^ ion from the pocket (see Figure 8b,d). Glu354/733 is the only CRD residue forming noticeable contacts with the mannose moiety, via OH2 and OH3. This state is much less stable than the others (see the ligand/CRD binding affinities listed in Figure 7b). Finally, *state C* was only observed for the mannose binding to the MRC1 CRD. In this case, the mannose OH3 and OH4 form contacts with the Ca^2+^ ion, while OH2 interacts with Glu733. Following a shift of the mannose moiety, Glu725 no longer interacts with the ligand. Instead, the OH6 group now forms hydrogen bonds with Glu737 (see Figure 8e). This state is the most stable one (see Figure 7b) and is specific to the MRC1 CRD, even though the DC-SIGN CRD also has Glu358 in the same position as MRC1 Glu737. The mannose loss of contacts with Glu725 is however compensated by the OH3 group forming contact with Tyr729 (see Figure 8e), which is not possible in the DC-SIGN CRD, where a valine residue occupies the same position. *State C* notably contributes to the longer mannose binding times that we observed for the MRC1 CRD compared to the DC-SIGN system (see Figure 7a). Further exploration of the residues lining the glycan binding site show that, for the mannose group to bind in the *state C* orientation and form contacts with Glu358/737, Asn365/747 must rotate away from the Ca^2+^ ion (see Figure 9a). This rotated conformation (termed *far state*), which is characterized by an increase in the Ca^2+^/Asn-Nδ2 distance, appears to be more frequent in the MRC1 CRD than in DC-SIGN (see the distributions in Figures 9b and 9c). In particular, the distributions of the Ca^2+^/Asn-Nδ2 when considering only structures with mannose bound on the CRDs (Figure S6a) show that Asn365 never occupies the far state in DC-SIGN, meaning that this rotation only occurs in DC-SIGN after mannose unbinding (see Figure S7). In the simulation of the CRDs without ligand, Asn365/747 is initially in its close state (which correspond to the crystallographic structure) and stays so for the whole simulation in DC-SIGN, while it rapidly switches to the far state in MRC1 (see Figure S6b,e). We ran additional MD simulations of the CRDs without ligand (using the same parameters as in Table 1), but starting this time with Asn365/747 in its far state. For both CRDs, Asn365/747 stayed in the far state for the whole trajectory and never switched back to the close state (see Figure S6c,f). In the PDB, the C-type lectin domains crystallographic structures usually include the characteristic calcium atom in their glycan binding pocket. In that case, the asparagine residue corresponding to Asn365/Asn747 (from DC-SIGN and MRC1 respectively) is in its close state, with its oxygen atom forming a coordination bond with the ion. However, we could also find CRD structures that were crystallized without calcium, such as 1BUU^79^ and 1XAR,^80^ and whose glycan binding pocket Asn residue is in the far state, thus supporting the biological relevance of this conformation.

**Figure 7:**
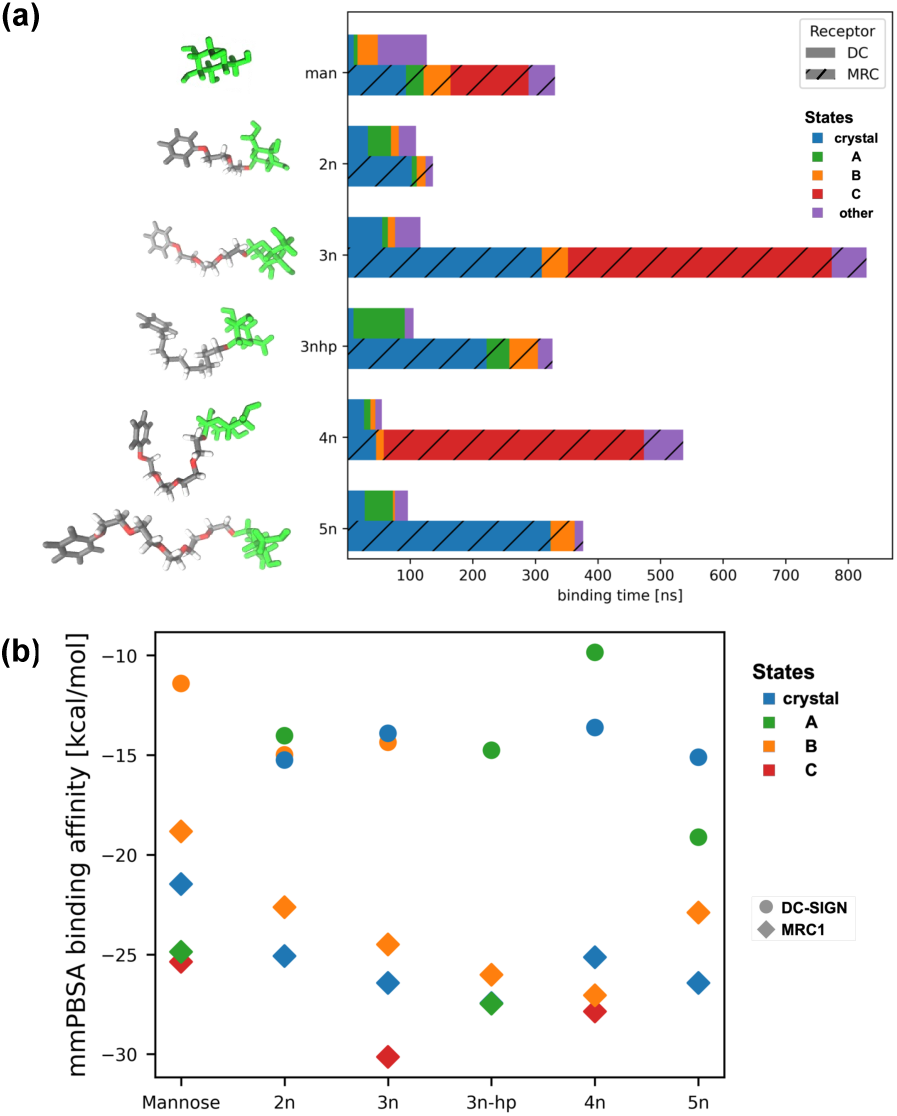
(a) Distribution of the mannose binding states for all the possible ligand/ CRD combinations (6*2 systems, the three replicas for each combination have been concatenated), blue: crystal position, green: A state, orange: B state, red: C state (MRC1 specific), lilac: other. ( b) MMPBSA binding affinities ( in kcal.mol^-1^) for the different ligand binding states on the DC-SIGN (circles) and MRC1 (diamonds) CRDs. Note that for the 3n-hp ligand with MRC1, the crystal position and state A have the same values, and the blue and green diamonds are superposed.

**Figure 8:**
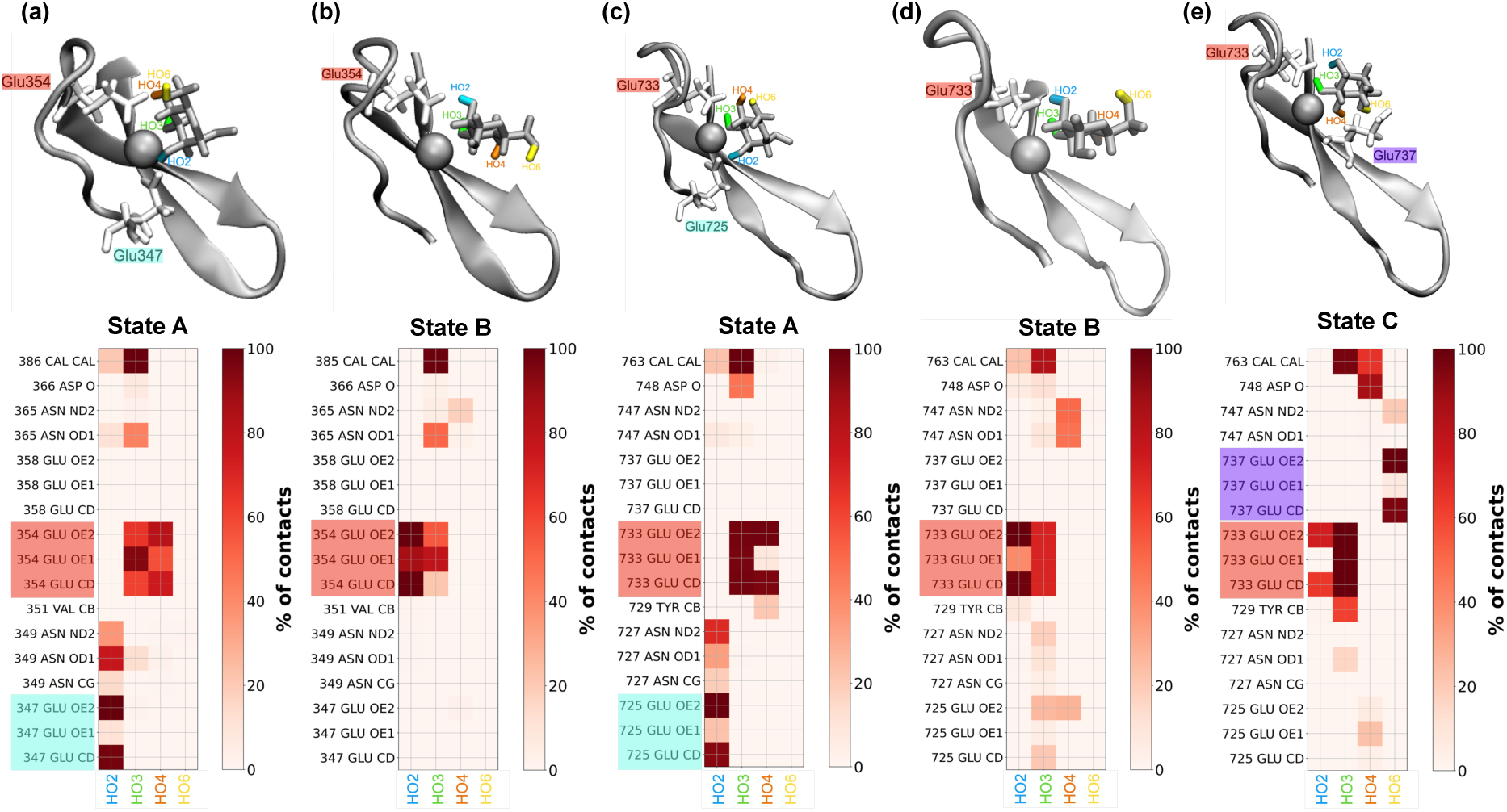
Mannose binding modes in the CRD glycan binding pockets with the associated contact maps. DC-SIGN: (a) state A, (b) state B. MRC1: (c) state A, (d) state B, (e) state C

**Figure 9:**
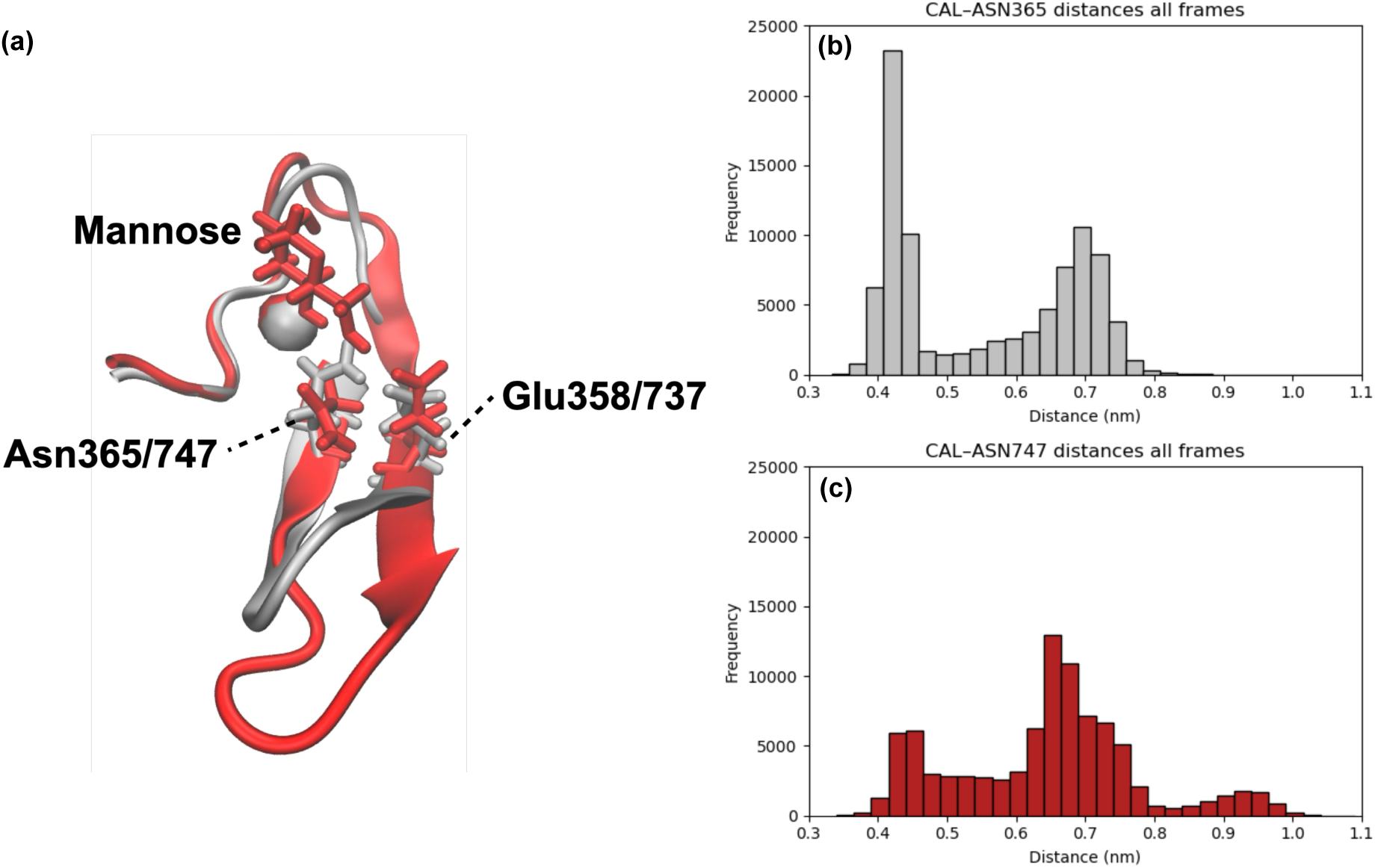
(a) Aligned structures of the glycan binding pockets for the DC-SIGN (in grey) and MRC1 (in red) CRDs with the mannose positioned in the C binding state. DC-SIGN Asn365 (in grey) is close to the Ca^2+^ ion, while MRC1 Asn747 (in red) has rotated away. (b) Distribution of the Ca^2+^-Nδ2 distance for Asn365 in DC-SIGN for all trajectories (with and without bound mannose). (c) Distribution of the Ca^2+^-Nδ2 distance for Asn747 in MRC1 for all trajectories (with and without bound mannose).

### Alternative binding sites on the CRDs surface

Previous experimental studies using NMR spectroscopy reported secondary binding sites on C-type lectins,^28, 81^ and even showed that for DC-SIGN, ligands binding these secondary sites might allosterically enhance glycan recognition by the main glycan binding pocket.^82^ During our MD simulations, we also observed short-lived (lasting for 5-10 ns) rebinding events of the mannose-based ligands (but not for simple mannose) after leaving the glycan binding pocket (see Table S1 for a complete summary of the binding residues in the secondary binding sites). These rebinding events are also visible in the ligand RMSD plots shown in Figure S3, as they result in a stabilisation of the RMSD fluctuations after unbinding from the main glycan binding site. This is particularly the case for the 3n-hp ligand, as its hydrophobic linker favors rebinding on the protein’s surface instead of solvation in water.

While earlier studies on alternative sites in DC-SIGN are mostly related to aromatic biphenyl substrates like the quinolone scaffold,^83^ we could observe a similar phenomenon with only a simple linear linker and a benzene ring. For the simulations with the DC-SIGN CRD, the ligand binds on several occasions next to Phe313 (Figure 10a, in purple), which has been proposed in literature to be part of an extended canonical binding site (CBS) for large enough ligands.^26, 28^ Two others secondary binding sites are located near the 𝛼_2_-helix of the lectin fold. One directly covers the helix (in blue on Figure 10a) and concurs with the site II from ref. 81, while the second one (in orange on Figure 10a) lies below the 𝛼_2_-helix. The green site on Figure 10a lies next to the 𝛼_1_-helix, with the ligand forming contacts with Trp277 and His278, and can be related to site V from ref. 81. Finally, we also observed frequent interactions between the ligands and the 𝛼_1_-helix lower part, on one side, and the protein’s own termini on the other side (in red in Figure 10a), but, as far as we know, these were not previously reported in NMR studies.

**Figure 10:**
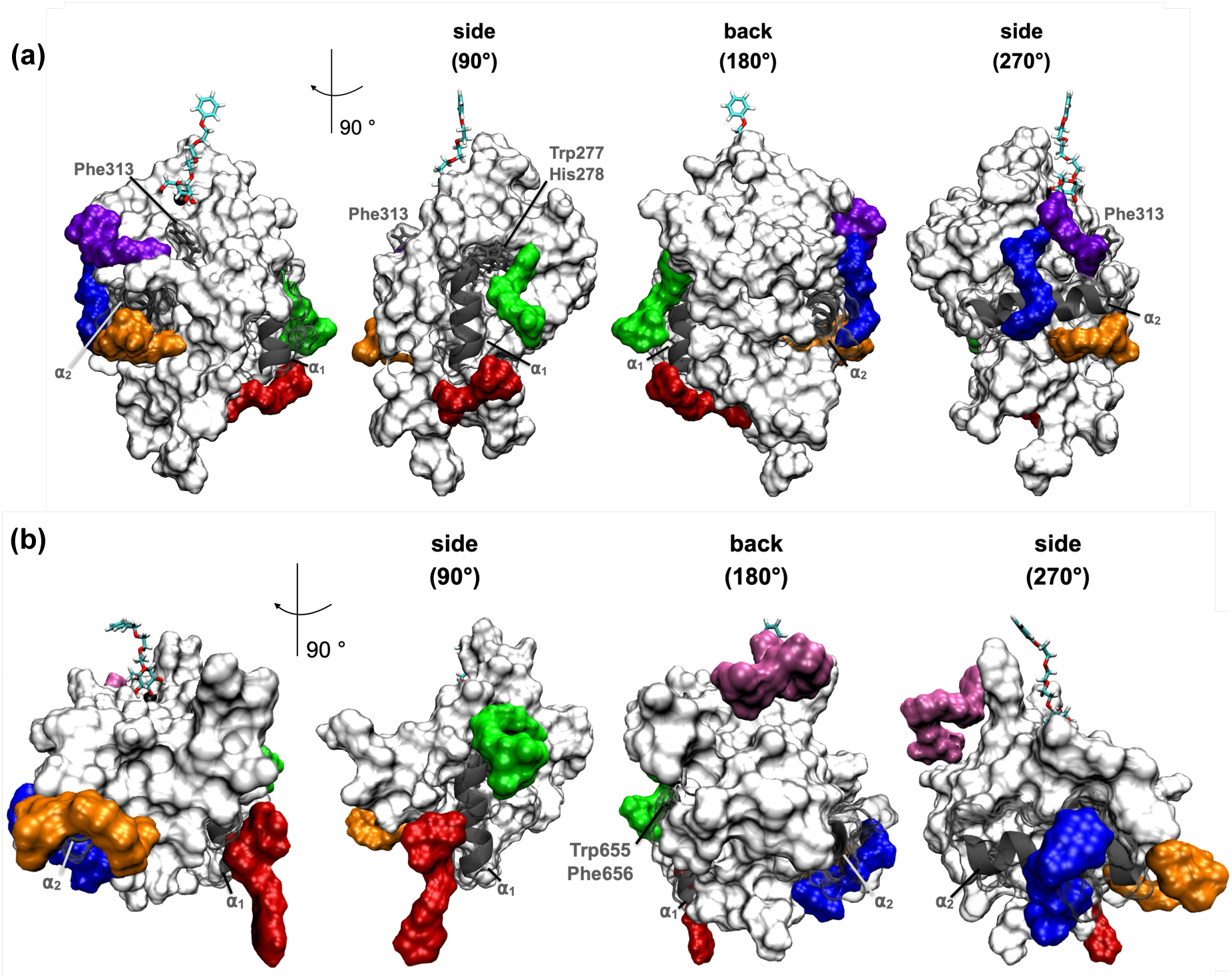
Alternative binding sites observed for both CRDs. The CRD is shown from four different angles in white surface representation with the ligands binding alternative sites in various colors. The 3n ligand bound to the canonical binding pocket is shown in licorice representation. Secondary elements and residues important for ligand binding are highlighted in grey (a) DC-SIGN, (b) MRC1.

Simulations of the MRC1 CRD also led us to observe secondary ligand binding events. Like in DC-SIGN, the ligands bind next to the 𝛼_1_-(in green and red on Figure 10b) and 𝛼_2_-helices (in blue and orange on Figure 10b). The green binding site around Trp655 and Phe656 can be related to the one observed next to Trp277 and His278 in DC-SIGN, as both residue pairs occupy the same position in the CRDs sequences (see the alignment in Figure S8). Ligand binding behind the 𝛼_2_-helix (in orange on Figure 10b, 270° orientation) seems to be specific to the MRC1 CRD. Additionally, we could observe ligand binding below the CBS long loop (in mauve on Figure 10b). In contrast to DC-SIGN, there are no calcium atoms located in this pocket (see Figure 2b). The binding site at Phe313 of DC-SIGN is not matched in MRC1, as corresponding Phe695 residue in the MRC1 sequence (see Figure S8) points inwards, towards the core of the protein, and thus does not form a convincing site of interaction for the ligands.

## 4. Conclusion

We used all-atom and coarse-grain simulations on the classical level to extensively characterize the carbohydrate recognition domain of two mannose receptors, DC-SIGN and MRC1, and to investigate the binding modes of various mannose based ligands on these CRDs. For both receptors, the simulations highlight the low stability of the ligand in the glycan binding pockets. Starting from the crystallographic position, the reorientation of the mannose group during the MD trajectories leads to three additional binding states that involve different residues on the CRDs surface. In particular, the binding of mannose in *state C*, which is specific to the MRC1 CRD, requires the rotation of an asparagine residue (Asn365/747). This side-chain reorientation does not happen (or happens only after mannose unbinding) in the DC-SIGN CRD. The MD trajectories also show secondary ligand binding events on alternative binding sites on the CRDs surface. The higher structural stability of the MRC1 glycan binding pocket and the possibility of mannose binding via *state C* result in a higher mannose affinity for MRC1 compared to DC-SIGN, which questions DC-SIGN’s suitability as a target in retinoblastoma photodynamic therapy.

However, both receptors present a more complex structure in the cell that might drastically change the results obtained when working on the CRD alone. Indeed, DC-SIGN assembles as a homotetramer^84^ (see Figure 1a) and is thus able to simultaneously bind several mannose groups. Furthermore, recent experimental work by Porkolab et al.^85^ highlight an unbinding/rebinding mechanism, where the ligand sequentially binds on the different CRDs, which is likely to enhance its affinity for DC-SIGN. In addition, some of the alternative binding sites that were identified on the surface of the DC-SIGN CRD monomer might no longer be accessible in the homotetramer. On the other hand, the MRC1 CRD is inserted in fourth position, within a linear chain of C-type lectin-like domains (see Figure 1a). The CRD4 domain is surrounded by CRD3, which cannot bind mannose, and the C-type CRD5, which is missing the equivalent of Tyr729 and Glu737, thus making the possibility of mannose binding in *state C* impossible.

In this perspective, further work will address the multimeric structures of DC-SIGN and MRC1 by modeling the oligomeric assemblies of several CRDs for both receptors and investigate how this global description impacts the mannose binding mechanism in both systems. On the longer term, we will build on the knowledge acquired regarding the mannose/glycan binding site interface in both CRDs and complete receptors to design new glycomimetic ligands with an enhanced affinity for DC-SIGN compared to MRC1.

## Acknowledgments

This work was supported by the ANR (SIMPA-ANR-23-CE29-0022) and by the “Initiative d’Excellence” program from the French State (Grant “DYNAMO”, ANR-11-LABX-0011-01). Simulations were performed using the HPC resources from LBT/HPC and GENCI/TGCC (Grant A018071713)

## Supporting Information Available

Abbreviation list, additional data regarding the force-field choice, coordinates covariance the CRDs, ligand and calcium RMSD plots and information regarding the ligand alternative binding sites are available as a pdf file. All MD trajectories (without solvent) and their topologies for each simulation are deposited in Zenodo (https://zenodo.org/records/17550324). All the analysis scripts are available in a public github repository (https://github.com/sinageissler/DC-MRC1-paper1).

