## Supplementary Information for "A comparative investigation of the mannose binding interface in DC-SIGN and MRC1 carbohydrate recognition domains with all-atom molecular dynamics simulations"

### List of acronyms and abbreviations (in order of appearance in the manuscript)

PDT: Photodynamic therapy

PS: Photosensitizer

MR: Mannose receptor

CTLD: C-type lectin-like domain

CRD: Carbohydrate recognition domain

DC-SIGN: dendritic cell-specific ICAM-3-grabbing non-integrin

Rb: Retinoblastoma

MRC1: Mannose receptor C-type 1

ITC: Isothermal titration calorimetry

MD: Molecular dynamics

PDB: Protein data bank

VMD: Visual molecular dynamics

RMSD: Root mean square deviations

MM/PBSA: Molecular mechanics/Poisson-Boltzmann surface area

RMSF: Root mean square fluctuations

BD: Brownian dynamics

CBS: Canonical binding site.

### Additional comment regarding the choice of the force-field for the simulations.

We chose to use the CHARMM36m (Huang et al., 2017) force-field with TIP3P water (Onufriev & Izadi, 2018) as it is well tested and widely used for protein simulations in the molecular modeling community. Before production, a test-system (DC-SIGN, calcium and N-Acetylglucosamine) was used to determine the best suited calcium model for this setup as overbinding of ions (in particular multivalent ones) is a known problem with the CHARMM36m forcefield(Ahmed, Papaleo, & Lindorff-Larsen, 2018). When using either a multisite calcium model (Zhang, Yu, Liu, & Song, 2020) or rescaling of the calcium atom's charge (Duboue-Dijon, Javanainen, Delcroix, Jungwirth, & Martinez-Seara, 2020) , the $Ca^{2+}$ ion needed for ligand binding left the binding pocket during the simulation. As the goal of this study was to investigate the atomic interactions in ligand binding to the pocket, unbinding of the pocket calcium was undesirable in this setup, leading to the decision to use CHARMM36m's default calcium model for the simulations, while still keeping overbinding in mind. Nevertheless, *realistic* dynamics should be retained, which is why no position restraints were used in the production step of the simulations. This resulted in relatively quick unbinding of the ligand in all DC-SIGN simulations but not in all MRC1 simulations, providing valuable insights to the differences between the two receptors, which was the purpose of this study.

**Figure S1:** RMSD plots showing the equilibration of the protein structure during the whole procedure, with three equilibration steps of 125 ps, followed by three equilibration steps of 500 ps, and the beginning of the production step. Yellow lines: DC-SIGN replicas, Red lines: MRC1 replicas.

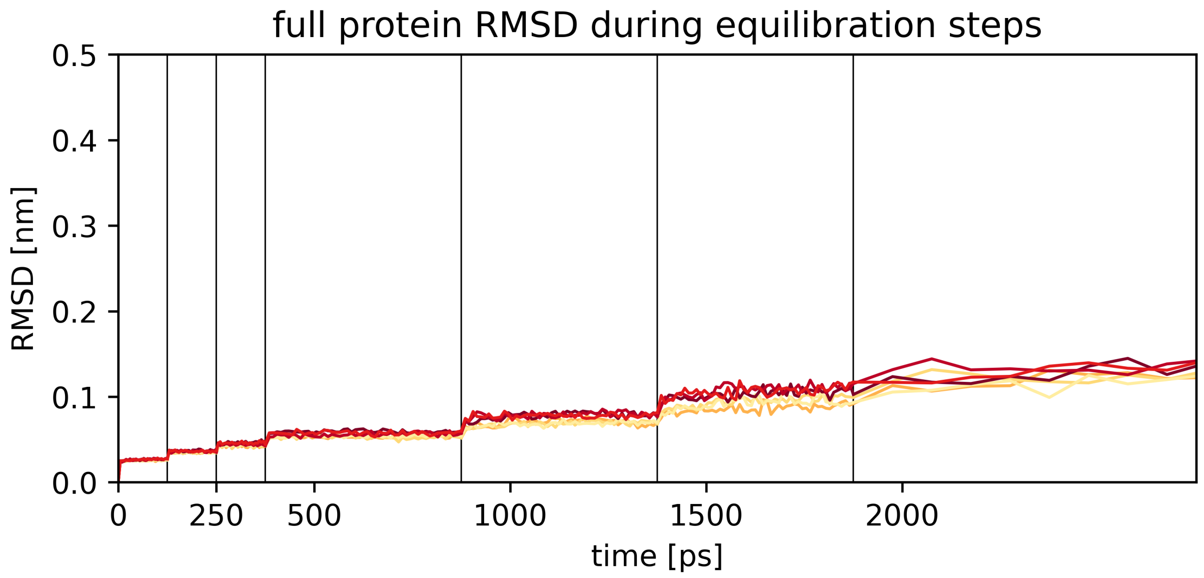

**Figure S2:** Principal components analysis for DC-SIGN and MRC1 CRDs in their apo state.

Cumulative variance for the ten first eigenvectors: (a) DC-SIGN, (b) MRC1

RMSF of each Cα along the five first eigenvectors, the glycan binding pocket is highlighted in grey: (c) DC-SIGN, (d) MRC1

RMSF of each Cα along eigenvectors 6-10, the glycan binding pocket is highlighted in grey: (e) DC-SIGN, (f) MRC1

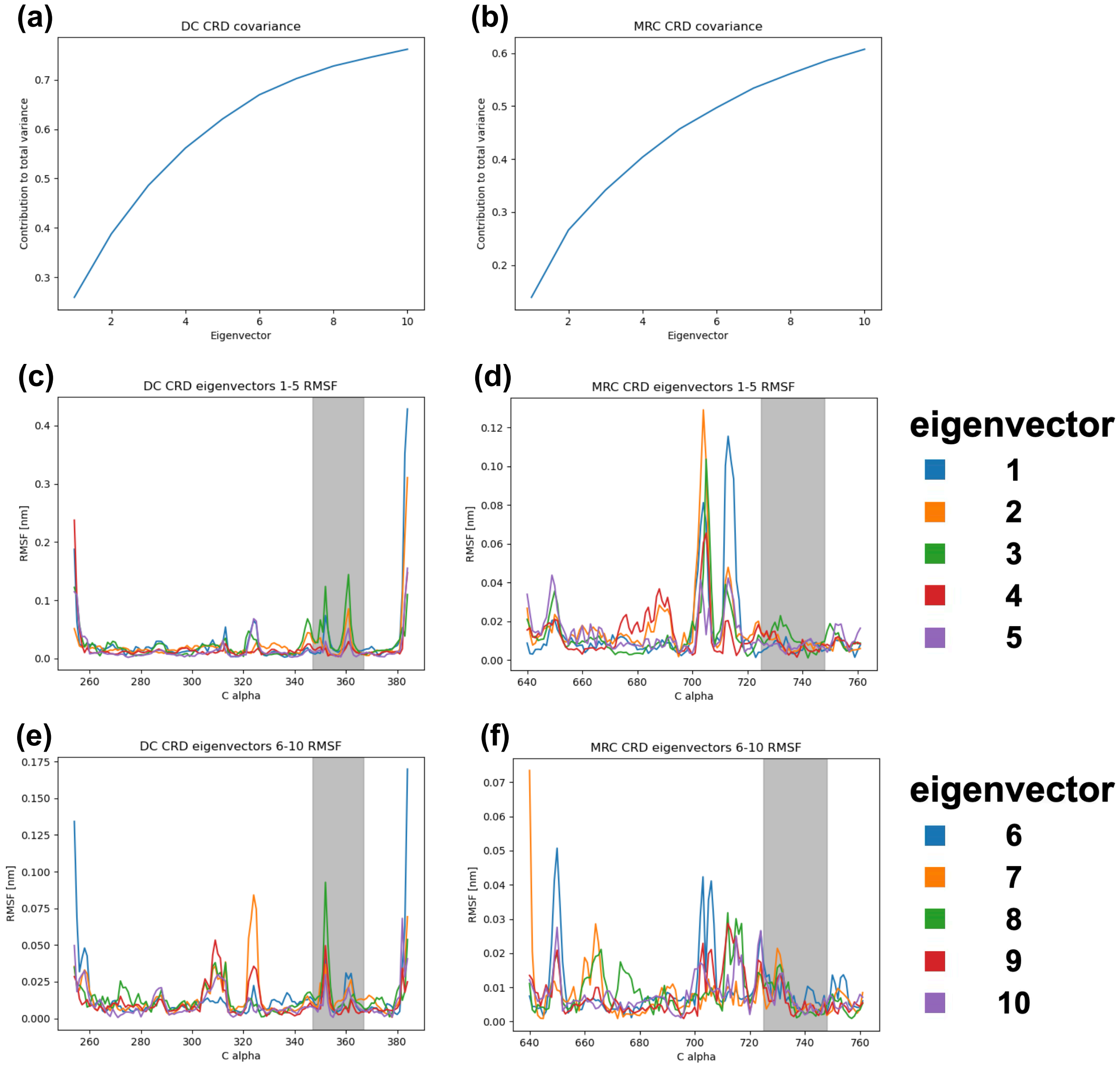

**Figure S3:** RMSD of the ligand (in green, smoothed values with a 20 frames sliding window) and $Ca^{2+}$ ion (in blue) with regards to the glycan binding site. (a) DC-SIGN, (b) MRC1

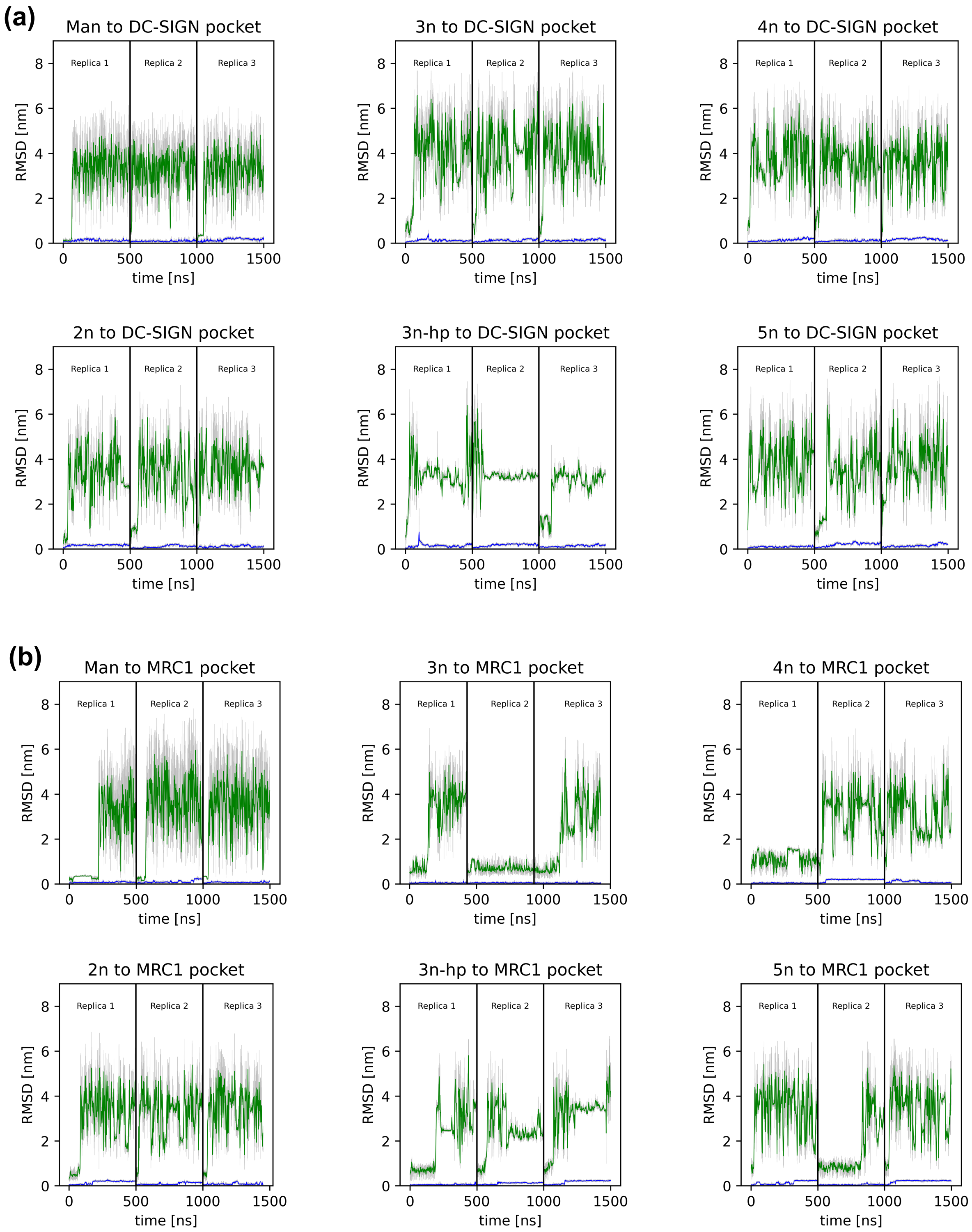

**Figure S4:** Distance between the glutamate residues surrounding the $Ca^{2+}$ ion in all the trajectories. (a) Glu347-Glu354 in DC-SIGN, (b) Glu 725-Glu733 in MRC1.

Black dots: apo-protein, single replica, Blue: mannose ligand, three replicas, Turquoise: 2n-ligand, three replicas, Green: 3n-ligand, three replicas, Yellow: 3n-hp-ligand, three replicas, Red: 4n-ligand, three replicas, Dark red: 5n-ligand, three replicas.

The background color changes highlight the three replicas for each system, while the dashed lines signal ligand unbinding events.

Distribution of the interglutamate distance over all the trajectories.

(c) Glu347-Glu354 in DC-SIGN, (d) Glu 725-Glu733 in MRC1.

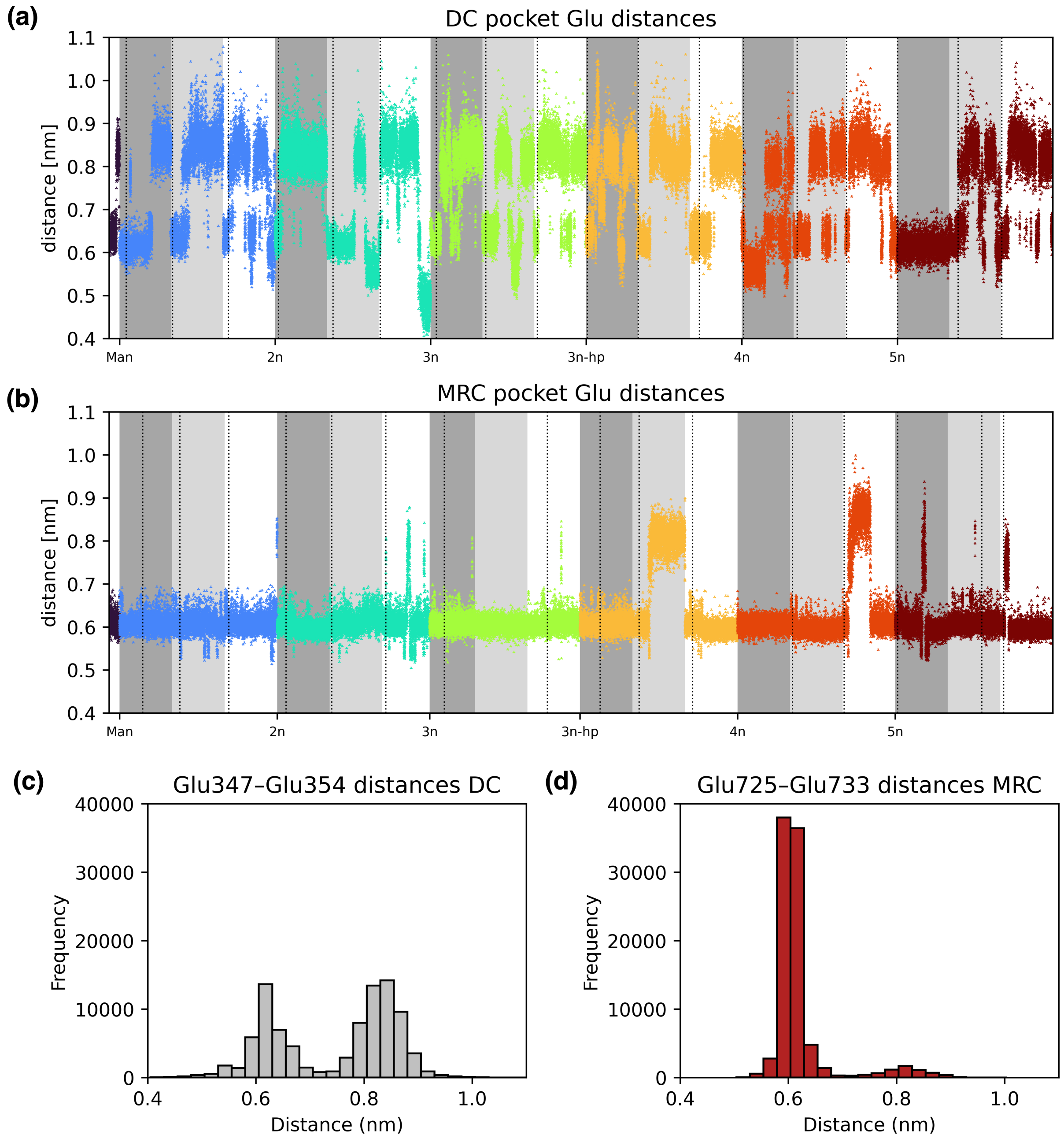

**Figure S5:** Snapshot of the 3n ligand bound on MRC1 CRD and interacting with Tyr729

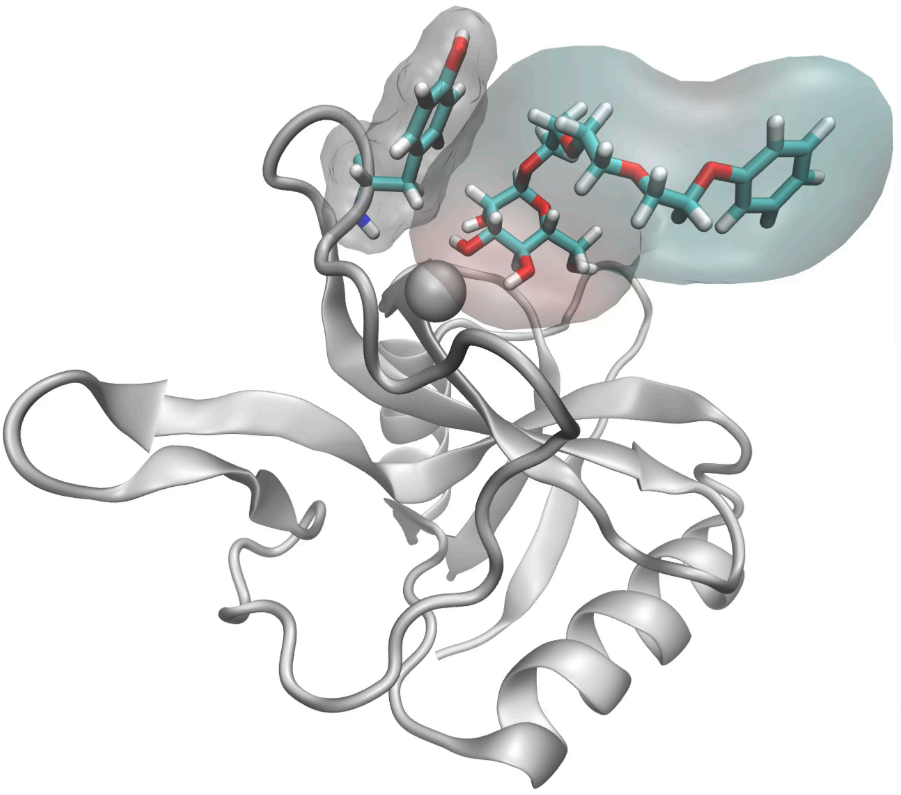

**Figure S6:** Distribution of the $Ca^{2+}$-sNδ2 distance for Asn365/747 in the DC-SIGN (in grey) and MRC1 (in red) CRDs.

DC-SIGN: (a) Mannose bound states in all trajectories, (b) apo-CRD starting from the crystallographic structure, (c) apo-CRD starting from a far orientation of Asn365.

MRC1: (d) Mannose bound states in all trajectories, (e) apo-CRD starting from the crystallographic structure, (f) apo-CRD starting from a far orientation of Asn747.

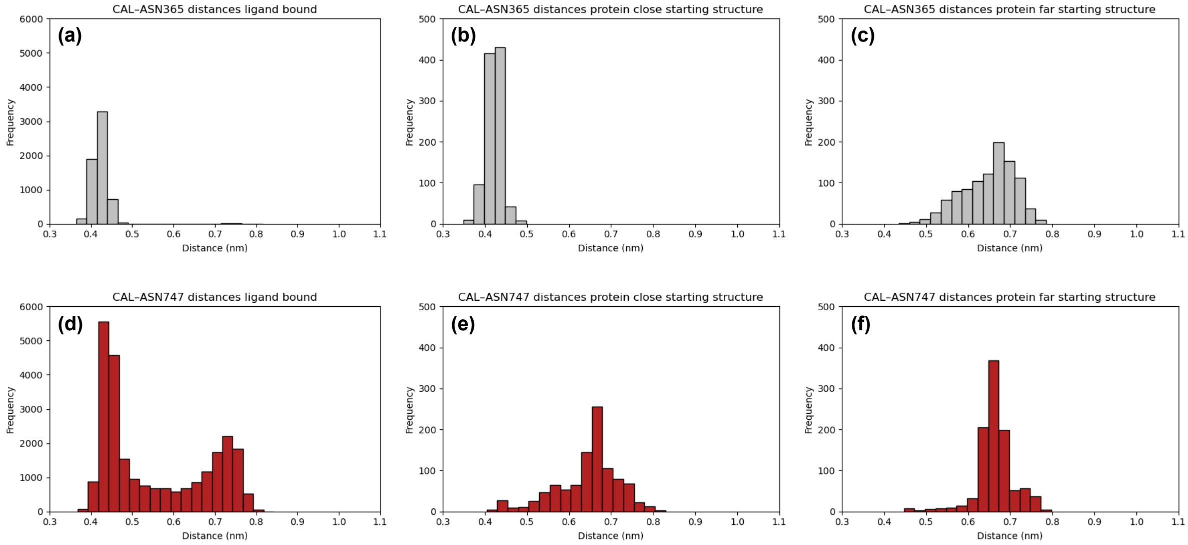

**Figure S7:** Time difference between ligand unbinding and Asn365/747 in the different replicas for all ligands. Grey bars: DC-SIGN, Red bars: MRC1. For DC-SIGN, ligand unbinding will always happen first.

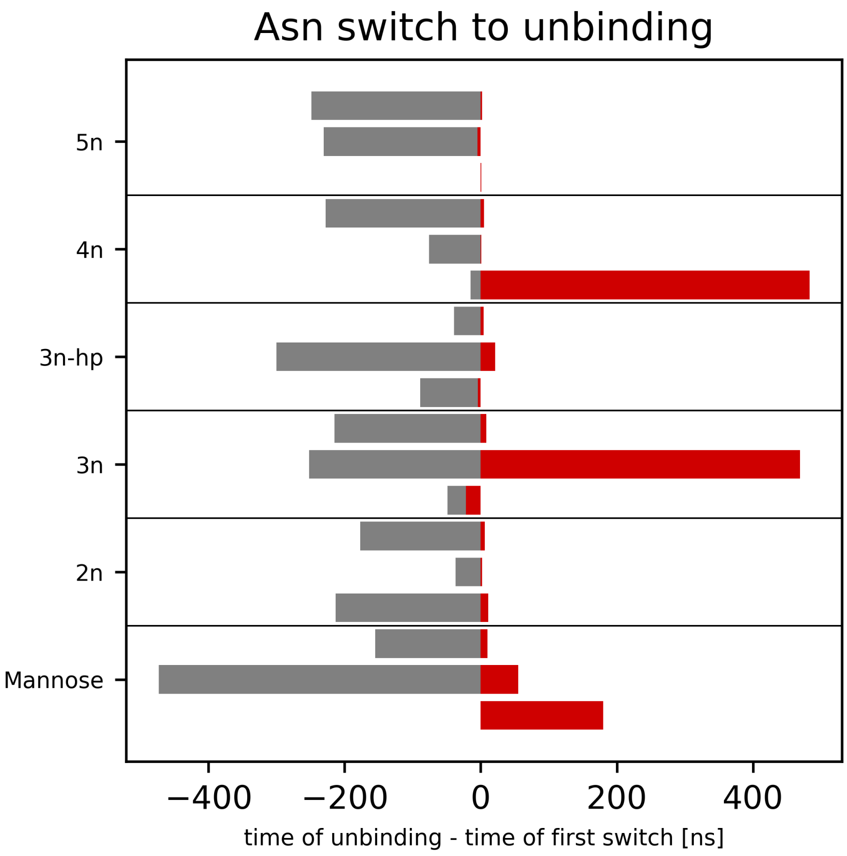

**Figure S8:** Sequence alignement of the DC-SIGN and MRC1 CRDs. Residues involved in the secondary ligands binding sites are highlighted in green and purple, matching the binding sites shown in Figure 10.

```
                                    277, 278 α1 helix                 α2 helix
DC  254     HPCPWEWTFFQGNCYFM----SNSQRNWHDSITACKEVGAQLVVIKSAEEQNFLQ---LQS   307
MRC 640     ------------LCFKLYAKGKHEKKTWFESRDFCRALGGDLASINNKEEQQTIWRLITAS   688
                         *: :    .:.::.*.:*   *: :*.:*. *:. ***: :      *
                                     655, 656

                 313                                                CBS
DC          SRSNRFTWMGLSDLNQEGTWQWVDGSPLLPSFKQYWNRGEPNNVGE-EDCAEFSG---N   362
MRC         GSYHKLFWLGLTYGSPSEGFTWSDGSPVS---YENWAYGEPNNYQNVEYCGELKGDPTM   744
            .  ::695:**:  . .  : * ****:     : *  *****  : * *.*:.*

DC          GWNDDKCNLAKFWICKKSAASC   384
MRC         SWNDINCEHLNNWICQI-----   761
            .*** :*:  : ***:
```

**Table S1 :** Summary of the secondary binding sites observed for the DC-SIGN and MRC1 CRDs.

| | | 2n | 3n | 3n-hp | 4n | 5n |
|---|---|---|---|---|---|---|
| **$\alpha_1$ top** | DC | ✓ | ✓ | ✗ | ✗ | ✓ |
| | MRC | ✓ | ✓ | ✓ | ✓ | ✓ |
| **$\alpha_1$ bottom / termini** | DC | ✓ | ✓ | ✗ | ✓ | ✓ |
| | MRC | ✓ | ✓ | ✓ | ✓ | ✓ |
| **$\alpha_2$** | DC | ✓ | ✓ | ✓ | ✓ | ✓ |
| | MRC | ✓ | ✗ | ✓ | ✓ | ✓ |
| **below $\alpha_2$** | DC | ✗ | ✗ | ✓ | ✓ | ✗ |
| | MRC | ✓ | ✗ | ✗ | ✓ | ✓ |
| **short loop** | DC | ✓ | ✗ | ✗ | ✗ | ✗ |
| | MRC | ✓ | ✗ | ✓ | ✗ | ✓ |
| **Phe313** | DC | ✗ | ✓ | ✗ | ✓ | ✓ |
